# Severe Acute Respiratory Syndrome Coronavirus-2 genome sequence variations relate to morbidity and mortality in Coronavirus Disease-19

**DOI:** 10.1101/2021.05.24.445374

**Authors:** Poonam Mehta, Saumya Sarkar, Ujjala Ghoshal, Ankita Pandey, Ratender Singh, Dharamveer Singh, Rahul Vishvkarma, Uday Chand Ghoshal, Ranjeet Maurya, Rajesh Pandey, Ravishankar Ramachandran, Punyasloke Bhadury, Tapas K Kundu, Singh Rajender

## Abstract

Outcome of infection with Severe Acute Respiratory Syndrome Coronavirus-2 (SARS-CoV-2) may depend on the host, virus or the host-virus interaction-related factors. Complete SARS-CoV-2 genome was sequenced using Illumina and Nanopore platforms from naso-/oro-pharyngeal ribonucleic acid (RNA) specimens from COVID-19 patients of varying severity and outcomes, including patients with mild upper respiratory symptoms (n=35), severe disease admitted to intensive care with respiratory and gastrointestinal symptoms (n=21), fatal COVID-19 outcome (n=17) and asymptomatic (n=42). Of a number of genome variants observed, p.16L>L (Nsp1), p.39C>C (Nsp3), p.57Q>H (ORF3a), p.71Y>Y (Membrane glycoprotein), p.194S>L (Nucleocapsid protein) were observed in similar frequencies in different patient subgroups. However, seventeen other variants were observed only in symptomatic patients with severe and fatal COVID-19. Out of the latter, one was in the 5’UTR (g.241C>T), eight were synonymous (p.14V>V and p.92L>L in Nsp1 protein, p.226D>D, p.253V>V, and p.305N>N in Nsp3, p.34G>G and p.79C>C in Nsp10 protein, p.789Y>Y in Spike protein), and eight were non-synonymous (p.106P>S, p.157V>F and p.159A>V in Nsp2, p.1197S>R and p.1198T>K in Nsp3, p.97A>V in RdRp, p.614D>G in Spike protein, p.13P>L in nucleocapsid). These were completely absent in the asymptomatic group. SARS-CoV-2 genome variations have a significant impact on COVID-19 presentation, severity and outcome.

## Introduction

Coronavirus Disease-19 (COVID-19) began in December 2019 in Wuhan City, the capital of Hubei province, China and led to the current pandemic that devastated the whole World. At the time of writing, 90,086,549 people in the world were infected with the Severe Acute Respiratory Syndrome Coronavirus-2 (SARS-CoV-2), of whom 1,934,939 (2.1%) had died. Interestingly, 85% of the subjects infected with the SARS-CoV-2 virus remain asymptomatic; of the remaining 15% symptomatic patients, about a third develop severe disease. Interestingly, both the morbidity and mortality rates differ across countries with mortality being as high as 15% in some countries and as low as 0% in others ^1^ https://coronavirus.jhu.edu/data/mortality (accessed on the 20^th^ Dec. 2020). The factors associated with variations in the outcome of SARS-CoV-2 infection remain largely unknown. In different geographical locations, variations in SARS-CoV-2 genome have been shown previously. Whereas the “A” (the ancestral type such as the bat coronavirus) and “C” are prevalent in the United States of America and Europe, the “B” subtype has been shown to be prevalent in East Asia ^2,3^

Earlier, we proposed the following factors that might predict the severity and outcome of SARS-CoV-2 infection; (A) host factors ^4,5^: (i) age of the patients, (ii) presence of co-morbid illness, (iii) variation in the host’s immune response to the virus due to difference in degree of T regulatory response (hygiene hypothesis), (iv) Bacillus Calmette-Guerin (BCG) vaccination, (v) variation in the gut microbiota ^6,7^; (B) agent (viral) factors: (i) viral load, and (ii) viral genome sequences. In our earlier study, we found that there was no difference in viral load among patients with and without gastrointestinal (GI) symptoms, the presence of which was independently associated with severe COVID-19 ^5^. Some studies focusing on the viral genome variations have reported that the variability in the symptoms could be linked to a few variations in the viral genome ^8,9^. However, such studies are based on secondary analysis of data obtained from Global Initiative on Sharing Avian Influenza Data (GISAID) database ^10^, which is prone to variations in the reporting of symptoms, their type and degree and the eventual outcome of the disease.

Among factors that could affect COVID-19 presentation, severity and eventual outcome, viral genome variations remain one of the most prominent and interesting factors. We undertook the whole genome sequencing of SARS-CoV-2 in nasopharyngeal swab samples from these patients and compared it with the reference genome sequence to investigate the relationship between SARS-CoV-2 genome variations and COVID-19 severity. We found that while some variations were shared across these classes, a number of mutations were more frequent in the symptomatic class and were completely absent in the specimens obtained from asymptomatic patients.

## Methods

### Patients

The experimental protocols were approved by the Institutional Ethics Committee of Sanjay Gandhi Postgraduate Institute of Medical Sciences. All methods were carried out in accordance with relevant guidelines and regulations. Informed consents were obtained from all participants. The samples were obtained from the patients diagnosed suffering from COVID-19 in a referral laboratory of a university hospital in northern India and from the patients admitted with the disease to the same hospital. All samples were collected between May and August, 2020. We undertook genome sequencing from naso-, oro-pharyngeal swab samples on a total of 115 COVID-19 patients with fatal disease (n=17), severe disease with gastrointestinal symptoms (GIS; n=21), mild disease with nasopharyngeal symptoms (NS; n=35), and asymptomatic infection (AS; n=42). The severity of the COVID-19 was assessed as described previously—(i) critical (required ventilator), (ii) severe (needed oxygen), (iii) moderate (although pneumonia was present, the patient did not require oxygen), and (iv) mild (only upper respiratory symptoms) ^11^. The gastrointestinal symptoms included anorexia, nausea, vomiting, abdominal pain and diarrhea.

### RNA isolation and molecular diagnosis of COVID-19

RNA was isolated using QIAamp Viral RNA kit (Qiagen, Hilden, Germany) using mini spin procedure according to the manufacturer’s instructions. RNA was quantified using nanodrop, followed by conversion of 50ng RNA into cDNA using high-capacity cDNA synthesis kit (Applied Biosystems, Foster City, California, USA). The first strand of cDNA was synthesized using random hexamers. SARS-CoV-2 detection was performed by quantitative real time PCR using Taqpath COVID-19 combo kit (Applied Biosystems, ThermoFisher Scientific, Foster City, California, USA). The kit uses spike (S) and nucleocapsid (N) protein regions for detection, which offer higher specificity due to lower risk of mutation in these regions.

### PCR amplification of SARS-CoV-2 genome

Samples with Cycle threshold (Ct) values of less than 28 were used for genome sequencing. cDNA synthesized in the above step was used for amplification of the SARS-CoV-2 genome using primers recommended by ARTIC Webpage: https://github.com/artic-network/artic-ncov2019/blob/master/primer_schemes/nCoV-2019/V3/nCoV-2019.tsv. The primers were pooled in four groups for optimal performance (Table 1).

**Table 1.** Set of primer pools used for SARS-CoV-2 genome amplification.

| **Primer pool** | **ARTIC primer name** |
| --- | --- |
| A | 13,17,19,23,25,27,29,31,33,35,37,39,41,43,47,49,51,53,55,67,71,73,75,91,97 |
| B | 10,12,16,18,20,22,24,26,28,30,32,34,36,38,40,42,48,50,52,54,64,66,68,70,74,92,94,98 |
| A-1 | 1,3,5,7,9,11,15,21,45,57,59,61,63,65,69,77,79,81,83,85,87,89,93,95 |
| B-1 | 2,4,6,8,14,44,46,56,58,60,62,72,76,78,80,82,84,86,88,90,96 |

PCR amplification of SARS-CoV-2 genome was carried out using AmpliTaq Gold 360 Master Mix (Applied Biosystems, Foster City, California, USA) with reaction constituents as 1X 360 master mix, 10mM dNTPs, 10µM primer pool. PCR cycling was carried out using conditions-initial denaturation at 95°C for 10 mins; 35 cycles of denaturation at 95°C for 30 secs: annealing at 59°C for 5 mins; extension at 72°C for 45 secs and final extension at 72°C for 10 mins on ABI Veriti PCR machine (Applied Biosystems, Foster City, California, USA). This was followed by AMPure beads (Cat. No. A63881, AMPure XP, Beckman Coulter, Pasadena, California, USA) based purification of cDNA and quantification using Qubit dsDNA HS assay kit (Cat. No. Q32854, Invitrogen, Carlsbad, California, USA).

### Illumina library preparation and sequencing

Amplicons generated by PCR in 4 reactions were pooled together and purified using 0.1X AMPure beads. 100ng of the pooled amplicons was used for subsequent sequencing using Nextera DNA Flex Library prep kit (Cat. No. 20018705, Illumina, San Diego, California USA). Briefly, tagmentation of purified amplicons was done by the addition of amplicon tagment mix (ATM) and tagment DNA buffer, as per the manufacturer’s protocol (Illumina Inc, San Diego, California USA) with incubation at 55°C for 15 mins with heated lid on. Tagmentation was stopped by the addition of tagment stop buffer (TSB) with incubation at 37°C for 15 mins. Further unique indexes and adapters (i7 and i5 adapters) were added to the samples. Index adapters were used for PCR amplification at 68°C for 3 mins, 98°C for 3 mins and 5-cycles of 98°C for 45 secs, 62°C for 30 secs, 68°C for 2 mins; and 68°C for 1 min. The PCR products were purified using Agencourt AMPure XP beads (Pasadena, California, USA). The quantity of the sequencing-ready library was measured using Qubit dsDNA HS assay kit (Cat. No. Q32854, Invitrogen, Carlsbad, California, USA) and quality was checked by Agilent DNA HS kit (Cat. No. 5067-4626, Agilent Technologies, Santa Clara, California, USA). Illumina’s MiSeq platform was used for sequencing.

### Miseq data analysis

The raw reads generated on MiSeq were checked for quality using FASTQC (version 0.11.8, Babraham Institute, Cambridge, UK). Trimgalore application was used to trim the reads containing bad quality and a minimum length of 40 base pairs was set as the threshold for the reads. The reads were converted from bam to fastq using bam2fastq (version 2.29) to align to the SARS-CoV-2 genome. HISAT2 was used to map the reads to the viral genome. Consensus fasta sequence was generated using samtools (version 1.9) and BCFtools on SARS-CoV-2 aligned files. The variants in the samples were called using BCFtools and VarScan.

### Oxford Nanopore Technology library preparation

The amplicons not covered in the first sequencing on Miseq were re-amplified separately and subjected to sequencing using nanopore sequencing technology. After purification for size and concentration check, 1ul of the purified amplicons were run on DNA1000 Agilent bioanalyzer (Cat. No. 5067-1504, Agilent, Santa Clara, California, USA). Further, 50ng of the amplicons were taken for end prep with NEBNext ULTRA II End Repair/dA tailing module (Cat. No. E7546, New England Biolabs, Ipswich, Massachusetts, USA) followed by incubation at 20°C for 5 mins and 65°C for 5mins. Further, 1.5µl of the end prep DNA was taken for native barcode ligation (EXP-NBD104 and EXP-NBD114, ONT, Oxford, UK) using Blunt/TA ligase master mix (Cat. No. M0367, New England Biolabs, Ipswich, Massachusetts, USA). The ligation mix was incubated at room temperature for 15 mins followed by AMPure bead purification. The purified product was quantified using Qubit dsDNA HS assay kit and 50ng of the product was further taken for adaptor ligation using adapter mix II and quick T4 DNA ligase (Cat. No. E6056S, New England Biolabs, Ipswich, Massachusetts, USA) followed by 20 mins incubation at room temperature. After adaptor ligation, purification was done using AMPure beads and short fragment buffer. Library was quantified using qubit dsDNA HS assay kit (Invitrogen, Carlsbad, California, USA) and 15ng of the library was used for sequencing. Sequencing flow cell was primed using flow cell priming kit (EXP-FLP002), and sequencing was performed on MinION Mk1B platform (Oxford Nanopore Technologies, Oxford, UK).

### Analysis pipeline for nanopore sequencing data

The raw Fast5 and Fastq files generated in MinKnow were basecalled and demultiplexed using Guppy basecaller (Version v.4.0.15). The ARTIC pipeline was used for bioinformatics analysis, which involved read filtering, primer trimming, amplicon coverage normalisation, variant calling and consensus building (https://artic.network/ncov-2019/ncov2019-bioinformatics-sop.html).

### Molecular phylogeny

For molecular phylogeny analysis, 3168 full length SARS-CoV-2 genome sequences submitted from India were downloaded from the GISAID database (November, 2020). Representatives of our SARS-CoV-2 genome sequences (25 types) were considered as part of this analysis. The reference genome of the SARS-Cov-2 RNA virus (NC_045512) was also considered in this study. All the genome sequences (3181) were subsequently aligned in MAFFT (https://mafft.cbrc.jp/alignment/server/). Based on the alignment, only 3020 full length genome sequences were further considered for downstream analyses. Twenty-five representative sequences we generated were subjected to maximum likelihood (ML) phylogeny based on Juke-Cantors model with bootstrap replicates (1000) in MEGA v7.0 ^12^. For the alignment containing 3020 full length genome sequences, nucleotide composition, codon usage, singletons and CpG were calculated in MEGA v7.0. Molecular phylogeny of the 3020 genome sequences were undertaken using NJ approach while taking into account the number of synonymous substitutions per synonymous site.

### Protein based annotation

Specific amino acid changes as a result of a mutation were annotated using SnpEff (version 4.5) ^13^. For this, the reference sequence of SARS-CoV-2 from Wuhan was used (NC_045512). For all proteins encoded by *ORF1ab* gene, the variants were annotated according to each mature protein. The analysis to figure out the conservation of these amino acids, multiple sequence alignment of the full-length sequences of six other coronaviruses was undertaken using Clustal Omega ^14^. The amino acid types were classified as per their type of side chains and polar nature.

## Results

### Clustering of mutations in the symptomatic class of patients

Some of the variants, such as p.16L>L (Nsp1), p.39C>C (Nsp3), p.57Q>H (ORF3a), p.71Y>Y (Membrane glycoprotein), p.194S>L (Nucleocapsid protein) were observed in similar frequencies across all categories of patients (Table 2). There were seventeen other variants which were observed only in the symptomatic class of patients (with gastrointestinal or nasopharyngeal symptoms or those who had succumbed to COVID-19). Out of the latter, one was in the 5’UTR (g.241C>T), eight were synonymous (p.14V>V and p.92L>L in Nsp1 protein, p.226D>D, p.253V>V, and p.305N>N in Nsp3, p.34G>G and p.79C>C in Nsp10 protein, p.789Y>Y in Spike protein), and eight were non-synonymous (p.106P>S, p.157V>F and p.159A>V in Nsp2, p.1197S>R and p.1198T>K in Nsp3, p.97A>V in RdRp, p.614D>G in Spike protein, p.13P>L in nucleocapsid). These mutations were completely absent in the asymptomatic category of the patients. However, the occurrence and frequency of these mutations did not differ significantly across three categories (dead, GI, NS) of the symptomatic patients.

**Table 2.** List of sequence variations observed in more than one instance.

| Sequence with change | Ge-<br>nomic<br>position | Coding<br>position | Location | Amino<br>acid<br>change | Struc-<br>ture<br>loca-<br>tion | Frequency n (%) |  |  |  |
| --- | --- | --- | --- | --- | --- | --- | --- | --- | --- |
|  |  |  |  |  |  | GIS<br>(N =<br>21) | NS<br>(N = 35) | Dead<br>(N = 17) | AS<br>(N =<br>42) |
| ATCTAGGTTT/CGTC<br>CGGGTGT | g.241C><br>T | Non-<br>coding | 5'UTR | --- |  | 16<br>(76.2) | 29<br>(82.86) | 15<br>(88.24) | 42 (100) |
| AAACACACGTC/TCAA<br>CTCAGTT | g.307C><br>T | c.42C>T | Nsp1 | p.14V>V | Sheet |  |  | 3 (17.65) | 0 |
| ACGTCCAACCTC/TAGT<br>TTGCCTG | g.313C><br>T | c. 48C>T | Nsp1 | p.16L>L | Sheet | 8<br>(28.09<br>) | 12<br>(34.28) | 5 (29.41) | 16<br>(38.09) |
| TAGCAGAACTC/TGAA<br>GGCATT | g.541C><br>T | c.276C>T | Nsp1 | p.92L>L | Sheet | 2<br>(9.52) |  |  | 0 |
| GACTATTCAAC/TCAA<br>GGGTAA | g.1121C<br>>T | c.316C>T | Nsp2 | p.106P>S | Loop | 2<br>(9.52) |  |  | 0 |
| GGGCGATTTTG/TTTA<br>AAGCCAC | g.1274G<br>>T | c.469G>T | Nsp2 | p.157V>F | Loop | 2<br>(9.52) |  |  | 0 |
| TTTGTAAAGC/TCAC<br>TTGCGAA | g.1281C<br>>T | c.476C>T | Nsp2 | p.159A>V | Loop | 2<br>(9.52) |  |  | 0 |
| ATGAGAAGTGC/TTCT<br>GCCTATA | g.2836C<br>>T | c.117C>T | Nsp3 | p.39C>C | Loop | 3<br>(14.28<br>) | 6<br>(17.14) | 3 (17.65) | 6<br>(14.28) |
| AAAATGCAGAC/TATT<br>GTGAAG | g.3397C<br>>T | c.678C>T | Nsp3 | p.226D>D | Helix |  |  | 6 (35.29) | 0 |
| GAGGAGGTGTT/CGCA<br>GGAGCCT | g.3478T<br>>C | c.759T>C | Nsp3 | p.253V>V | Helix |  | 6<br>(17.14) | 4 (23.53) | 0 |
| CAAATGTTAAC/TAAA<br>GGTGAAG | g.3634C<br>>T | c.915C>T | Nsp3 | p.305N>N | Turn |  |  | 6 (35.29) | 0 |
| GTCTTTGGAGC/AACA<br>AAACCAG | g.6310C<br>>A | c.3591C><br>A | Nsp3 | p.1197S><br>R | Loop | 2<br>(9.52) | 5<br>(14.28) | 2 (11.76) | 0 |
| CTTTGGAGCAC/AAAA<br>ACCAGTT | g.6312C<br>>A | c.3593C><br>A | Nsp3 | p.1198T><br>K | Loop | 5<br>(23.8) | 5<br>(14.28) | 3 (17.65) | 0 |
| TAGCTAGTGGG/TGGA<br>CAACCAA | g.13126<br>G>T | c.102G>T | Nsp10 | p.34G>G | Loop |  | 6<br>(17.14) | 3 (17.65) | 0 |
| ACTGCCGTTGC/TCAC<br>ATAGATC | g.13261<br>C>T | c.237C>T | Nsp10 | p.79C>C | Helix | 2<br>(9.52) |  |  | 0 |
| CCAGCTGTTGC/TTAA<br>ACATGAC | g.13730<br>C>T | c.290C>T | RdRp/Nsp<br>12 | p.97A>V | Loop | 5<br>(23.8) | 6<br>(17.14) | 3 (17.65) | 0 |
| CTTTATCAGGG/ATGT<br>TAACTGC | g.23403<br>A>G | c.1841A><br>G | Spike<br>protein | p.614D>G | Loop | 16<br>(76.2) | 29<br>(82.86) | 13<br>(76.47) | 42 (100) |
| AACAAATTAC/TAAA<br>ACACCAC | g.23929<br>C>T | c.2367C><br>T | Spike<br>protein | p.789Y>Y | Loop | 5<br>(23.8) | 6<br>(17.14) | 3 (17.65) | 0 |
| CTGTTTTTCAG/TAGC<br>GCTTCCA | g.25563<br>G>T | c.171G>T | ORF3a<br>protein | p.57Q>H | Helix | 5<br>(23.8) | 6<br>(17.14) | 4 (23.53) | 9<br>(21.43) |
| CTGCTGTTTAC/TAGA<br>ATAAATT | g.26735<br>C>T | c.213C>T | Mem-<br>brane<br>glycopro-<br>tein | p.71Y>Y | Loop | 5<br>(23.8) | 6<br>(17.14) | 4 (23.53) | 7<br>(16.67) |
| CGAAATGCACC/TCCG<br>CATTACG | g.28311<br>C>T | c.38C>T | Nucleoca<br>psid<br>phosphopr<br>oteins | p.13P>L | Loop | 5<br>(23.8) | 6<br>(17.14) |  | 0 |
| CGCAACAGTTC/TAAG<br>AAGAAAT | g.28854<br>C>T | c.581C>T | Nucleoca<br>psid<br>phosphopr<br>oteins | p.194S>L | Loop | 5<br>(23.8) | 6<br>(17.14) |  | 7<br>(16.67) |

### Molecular phylogeny

Based on molecular phylogeny, the ML tree consisting of SARS-CoV-2 genomes we generated revealed two main clusters - Clade I and II (Fig. 1). The Clade I consisted of 21 genome sequence types while the Clade II consisted of the remaining 4 types. Divergent sub-clades were encountered in both the clades with strong bootstrap support.

**Fig 1.**
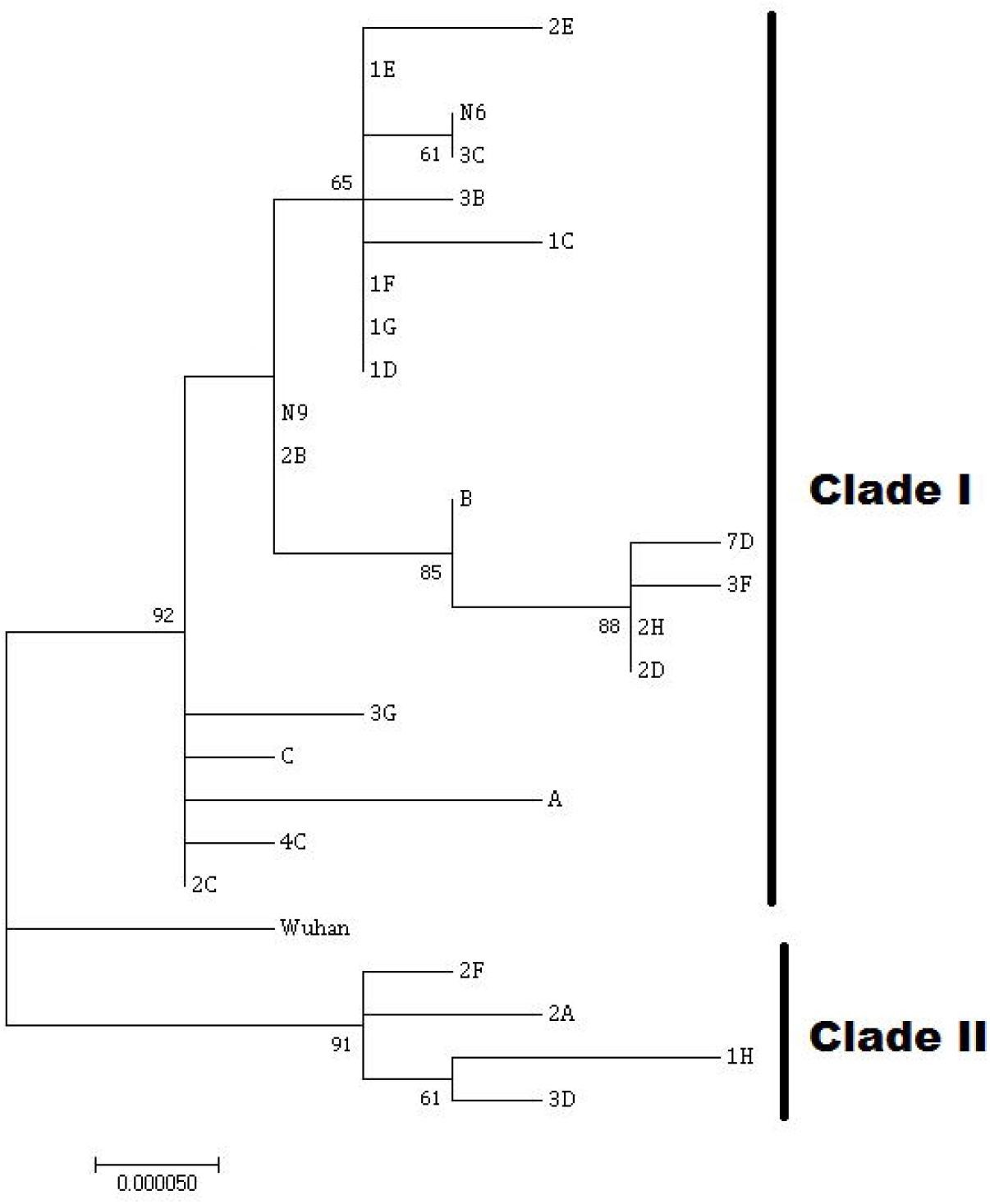
Maximum likelihood tree based on Juke-Cantors model for 25 representative SARS-CoV-2 genome types with bootstrap support (1000 replicates). The scale bar indicates 0.00005 substitutions per site.

In this study, the robust alignment consisted of 3020 full-length genome sequences with no ambiguity was undertaken. The relative synonymous codon usage (RSCU) and basic nucleotide composition (A%, U%, C%, and G%), AU and GC contents have been detailed in Supplementary Tables 1 and 2 respectively. Based on codon analysis, most of them tend to exhibit U ending. Interestingly, encountered CpGs were low (1640) based on the analysis of 3008 genome sequences. Interestingly, encountered CpGs were found to be low (1642) based on the analysis of 3020 genome sequences. From the genome alignment, 3157 singletons were identified. Based on the phylogeny of 3020 genome sequences, several clades were observed in the NJ tree (represented as condensed tree in Fig 2). Interestingly, a distinct divergent sub clade consisting of 2E genome type in our samples clustered with eight near complete SARS-CoV-2 genome sequenced from patients in Maharashtra and Gujarat, reflecting possible human connectivity during transmission.

**Fig 2.**
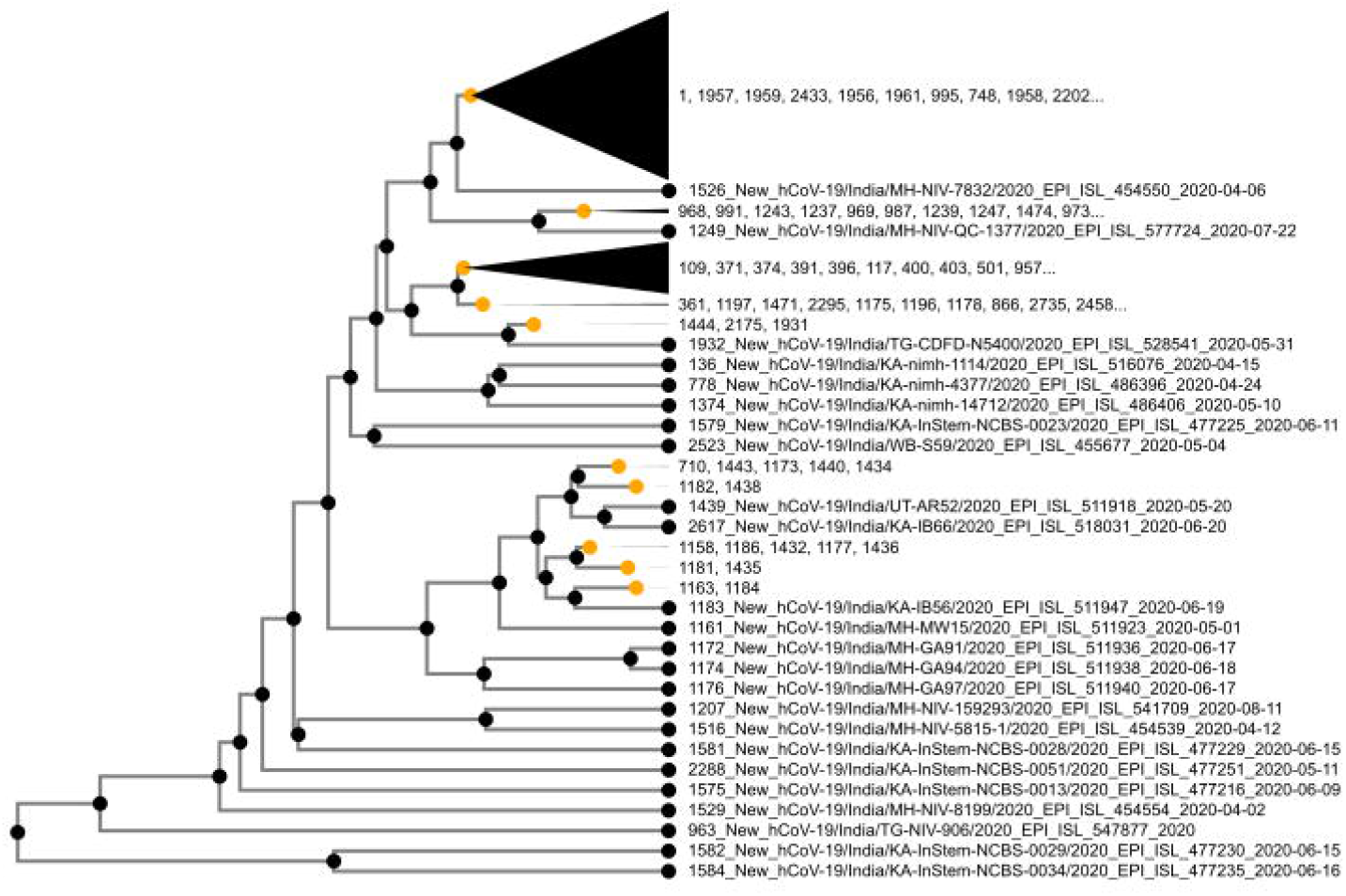
Condensed NJ tree of 3020 genome sequence dataset showing distinct nodes along with cladistic based on synonymous substitutions per synonymous site.

### Conservation of changes

Multiple alignment of viral protein sequences across the SARS, MERS and other virus families showed that the amino acid residues were conserved across the SARS family only (Fig 3). Most of the amino acid residues were not conserved in other viruses such as MERS. This showed significant changes in the protein sequences and the divergence of the SARS family from other viruses.

**Fig 3.**
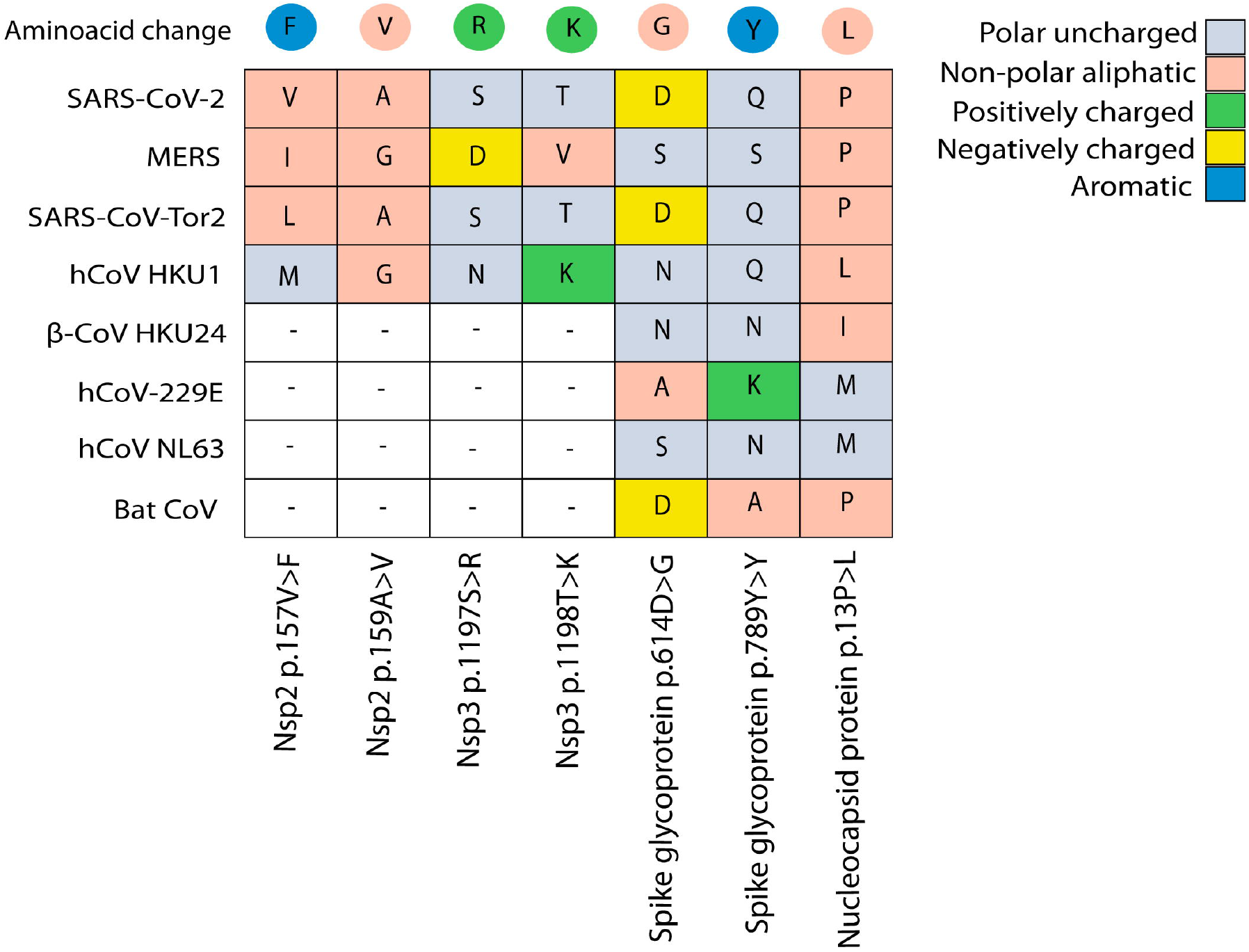
Protein sequence alignment showing conservation of amino acid residues across virus families.

### Structure-function analysis

We observed three mutations in the Nsp2 protein (C1121T, G1274T, and C1281T) in our samples (Table 2), which caused P106S, V157F, A159V amino acid substitutions, respectively. A159V changed into the same group of amino acid while V157F changed aliphatic to aromatic group. We mapped these positions in the full-length protein structure model (experimental structure not available). Out of two mutations present in the C-terminal loop region, V157F (valine to phenylalanine) may affect the structure-function while A159V (alanine to valine) may not affect protein function (Fig. 4). Also Nsp2 secondary structure composition suggests it to have large loop region between the N-terminal and the C-terminal, which may cause fluctuations in the structure.

**Fig 4.**
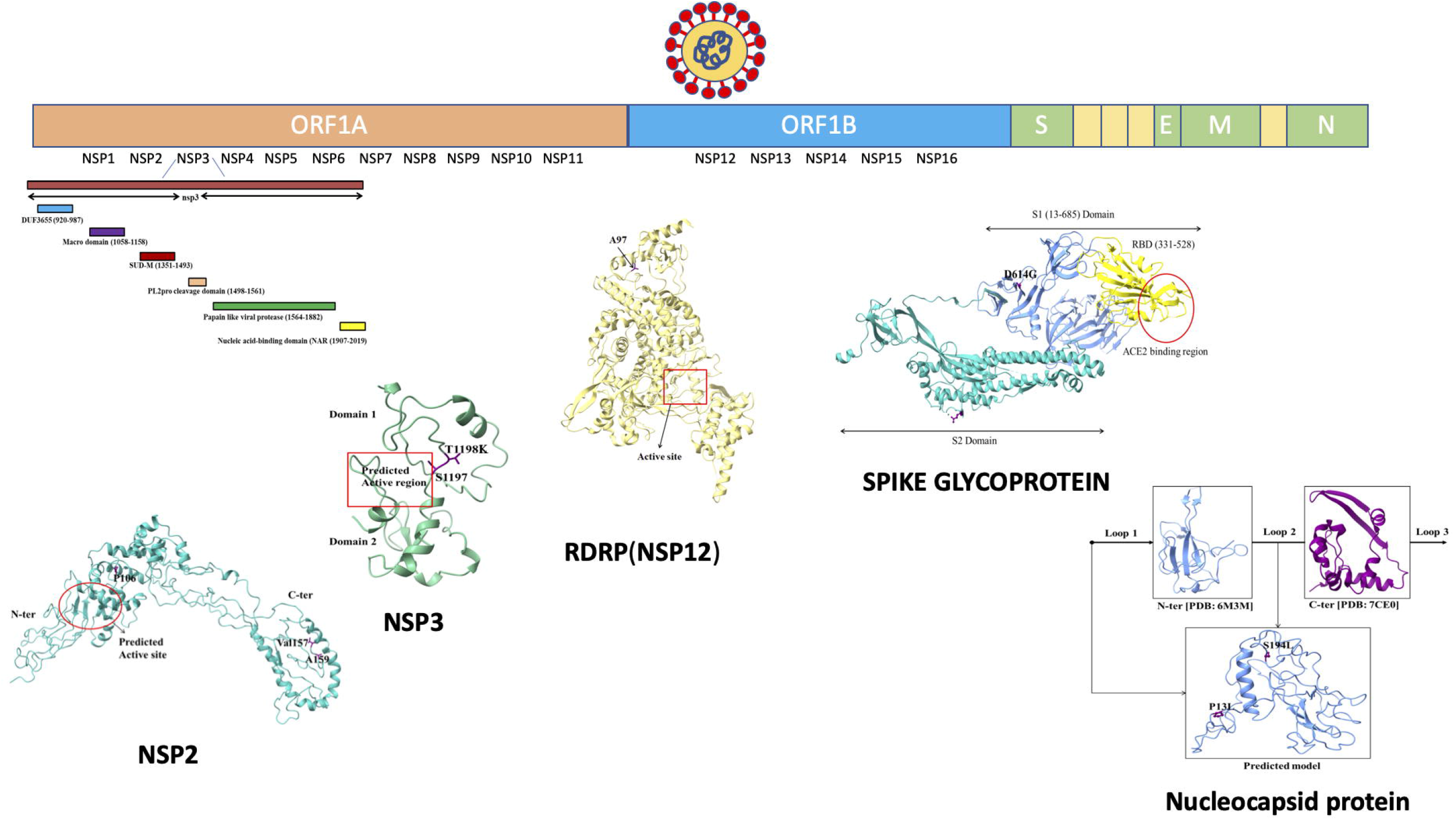
Structure-function analysis of non-synonymous mutations in the viral proteins. The position of the mutations corresponds to the gene position on the bar above.

Nsp3 has six conserved domains; DUF3655 (protein of unknown function) domain, macro domain (ADP ribose domain), single-stranded poly(A) binding (SUD-M) domain, PL2pro cleavage domain, PL-pro (Papain like viral protease) and nucleic acid binding (NAR) domain. The mutations in Nsp3 were seen in mainly 3 domains; DUF3655, NAR domain and ADP ribose domain [15]. DUF3655 and NAR domains do not have any experimental structure, but it is available for the ADP ribose domain [PDB: 6W6Y]. Therefore, we modelled the structure and mapped the location of the mutated residues. Two mutated residues (D226D and V253V) in the ADP ribose domain were present in the pocket and one (N305N) was present near the pocket (Fig. 4). The NAR domain had two domains with two mutations (S1197R and T1198K) present in the loop region of domain 1. Missense mutation T1198K substituted threonine (polar, neutral) with lysine (positively charged). The predicted active region is present between both the domains and both residues were present near the pocket. T1198K substitution is very likely to affect the structure-function of the protein. Both the mutations (S1197R and T1198K) in the Nsp3 protein lie in the nucleic acid binding domain and replaced polar uncharged amino acids with positively charged amino acids, suggesting enhanced interaction with viral genome due to the addition of a positively charged amino acid [16].

The RNA-directed RNA polymerase (RdRp) is responsible for replication and transcription of the viral RNA genome. The experimental structure is available with Nsp7 and Nsp8 complex, they activate and confer processivity to the RNA-synthesizing activity of the polymerase [17,18]. The mutation at position A97V has been previously shown to cause a change in secondary structure in RdRp enzyme [19]. The spike glycoprotein had an important point mutation at position D614G. Aspartic amino acid (D) is acidic and polar in nature while glycine is aliphatic and nonpolar. This mutation has been most widely investigated and affects spike protein conformation in a way to enhance the receptor binding ability of the spike protein [7,20].

ORF3a protein (ORF3a) [PDB: 6XDC] forms homotetrameric potassium sensitive ion channels (viroporin) and may modulate virus release [21]. It has a single mutation at Q57H, present in the loop helical region of the protein. Glutamine was substituted by Histidine, both are polar but different in nature, glutamine is neutral while histidine is basic in nature. This mutation might affect the function of the protein. The nucleocapsid (419aa) protein does not have full-length experimental structure available in the protein databank. Experimental structure is available for some regions only [PDB: 6M3M N-ter: (49-174) and PDB: 7CE0 C-ter: (255-366)]. Since these solved structural regions do not cover the mutations observed, we modeled the full length structure to map the position of mutated residues (P13L and S194L). P13L may not affect structure but S194L results in a different nature of amino acid, which may affect protein function (Fig 4). Both amino acids are present in the loop region.

## Discussion

In this study, we aimed at investigating the relationship between the SARS-CoV-2 genome sequence variations and COVID-19 severity. Among a number of mutations across the viral genome, a few clustered in symptomatic patients and were completely absent in asymptomatic patients. This included a 5’UTR mutation (g.241C>T), eight synonymous mutations (p.14V>V and p.92L>L in Nsp1 protein, p.226D>D, p.253V>V and p.305N>N in Nsp3, p.34G>G and p.79C>C in Nsp10 protein, p.789Y>Y in Spike protein), and eight non-synonymous mutations (p.106P>S, p.157V>F and p.159A>V in Nsp2, p.1197S>R and p.1198T>K in Nsp3, p.97A>V in RdRp, p.614D>G in Spike protein, p.13P>L in nucleocapsid). Sequence comparison with other coronaviruses showed a strong conservation of these amino acid residues within the SARS family of viruses, suggesting their critical role in protein function in this family. Some of the mutations we observed have been previously linked with disease severity on the basis of genome sequences downloaded from the public databases. For example, g.241C>T mutation in the 5’UTR, A97V in the RdRp protein, P13L in the nucleocapsid protein, D614G mutation in the spike protein, S1197R and T1198K in the Nsp3 protein were found more often in hospitalized patients/severe disease in comparison to those with mild disease ^9,15,16^. Comparison of the genotype frequency of five most important mutations across China, Italy, USA, UK and India showed significant changes in the frequency of alternate nucleotides at g.241C>T and p.614D>G positions (Fig 5).

**Fig 5.**
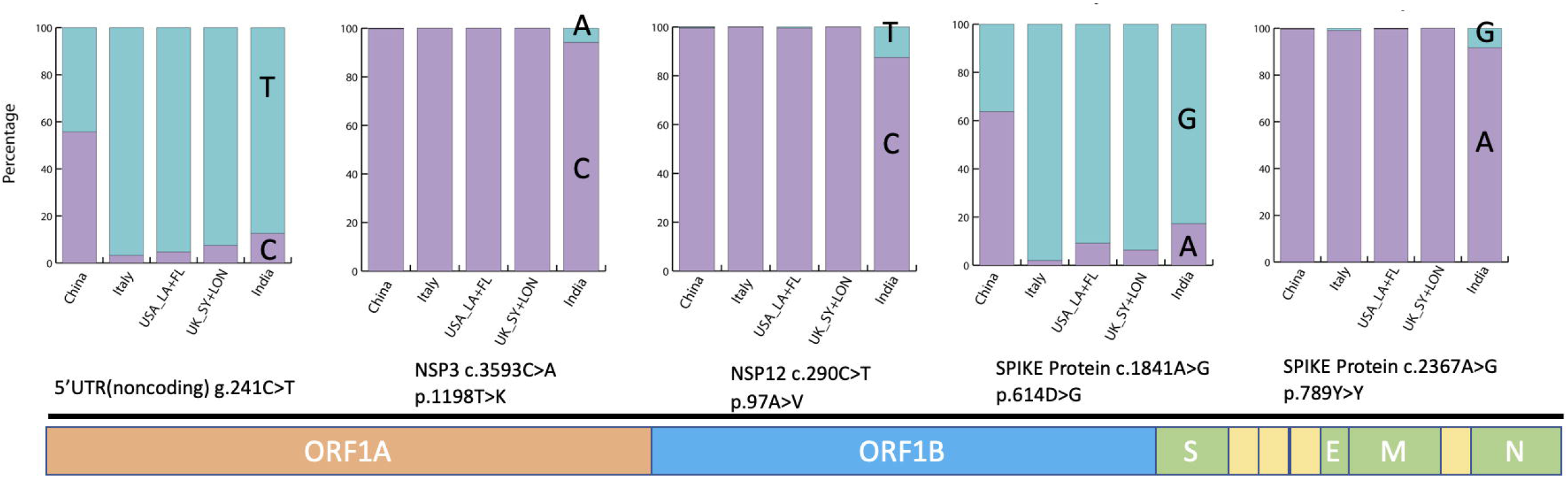
Comparisons of the frequencies of five most interesting mutations across China, Italy, United States (Los Angeles and Florida), UK (South Yorkshire and London) and India. This analysis was performed on data downloaded from the GISAID database.

Out of the mutations we observed in the symptomatic class only, D614G has been the most commonly discussed with respect to virus infectivity and virulence. The 614G form of the spike protein is more likely to assume an open conformation than the ancestral 614D form, resulting in improved binding to the ACE2 receptor. In human epithelial cells, the 614G variant replicated more efficiently than the 614D variant. Accordingly, D614G mutation has been found to increase transmissibility ^17,18^ and patients with 614G associated SARS-CoV-2 have been found to have higher viral load and show lower Ct values ^18^. Interestingly, in studies on hamsters despite variant infection ratios of upto 9 times in favour of 614D, 614G replicated to higher titres and outcompeted the D form and the 614G variant also seemed to be more stable in comparison to the ancestral 614D variant ^19^. However, 614G did not cause more severe disease than the ancestral 614D variant ^20^. We found that 614D is seen more commonly in the symptomatic patients with a more severe form of the disease, and was completely absent in the asymptomatic patients, though we did not find it to further differ as per severity in the symptomatic groups. Interestingly, in line with our observation, Volz et al. (2021) found that patients with 614G variant showed reduced odds of death, implying a less severe form of the disease, though this effect was not significant when adjusted for other known risk factors for severe COVID-19 outcome ^18^. Our results are in concordance with the initial observation of Volz et al, (2021) that 614G variant correlates with less severe forms of the disease. The presence of 614G form in asymptomatic patients could be behind faster spread of the virus after emergence of this mutant, for younger age of the infected patients and for less fatalities in the later months after its global spread from China/Europe. Higher frequency of 614G variant towards the end of the first COVID-19 wave could be behind the second wave characterized by a much faster spread than the first wave.

D614G has been the most controversial mutation with respect to its effect on virulence. While 614G has been established to result in higher infectivity ^19^, it has not been related to higher severity ^20^. At least three studies based on data from public databases suggest 614G to correlate with higher severity. Toyoshima et al, (2020) compared genomes from the GISAID database and found that 614G showed positive correlation with fatality ^21^. In another similar study, Biswas and Mudi (2020) found 614G to correlate with a more severe form of the disease ^9^. In yet another study, Nagy et al. (2021) downloaded genome sequences and classified the patients into mild outcome, hospitalized and severe outcome groups, the comparison of which showed increased severity with 614G ^16^. Nevertheless, all these studies are based on data from public databases and should not be taken as independent evidence supporting the correlation of 614G with a more severe disease. The reliability of the information provided in the database and further in-depth details including the types of symptoms, level of morbidity and the eventual mortality remain unreliable for conducting such studies with high confidence. In the instances where they have been compared, the conclusions regarding their impact on COVID-19 severity appear over-stated. We observed the 614D variant only in the symptomatic patients. Interestingly, a higher severity in the 614D variant has been supported by original investigations in the lone study with good sample size ^18^ and 614G has been ruled out for a higher severity ^20^.

A number of mutations we observed in the symptomatic patients were found to mildly or significantly affect protein function by in-silico assays. A97V has been previously shown to cause a change in secondary structure in RdRp enzyme ^22^. Both the mutations (S1197R and T1198K) in the Nsp3 protein lie in the nucleic acid binding domain and replaced polar uncharged amino acids with positively charged amino acids, suggesting enhanced interaction with viral genome due to the addition of a positively charged amino acid ^23^. However, functional assays are required to establish their effect on the viral survival, interaction with the host, entry inside the cells, immunogenicity and eventually the development of symptoms. Nevertheless, their clustering in the symptomatic group of patients is quite interesting and suggests their significant impact on host-virus interactions and immune response. Functionally important viral genome variations could have implications in herd immunity, drug action and the efficacy of vaccines. The impact of D614G variant on vaccine development has been tested; fortunately, 614G variant is equally sensitive to neutralization using serum samples ^19^. Based on the NJ phylogeny of 3020 genome sequences from India, the distinct subclade consisting of genome data represented by type 2E along with eight genome sequences from patients of Maharashtra and Gujarat reflected connectivity of transmission with possible signature of emerging divergence in these SARS-CoV-2 genomes. In India at an early stage, various SARS-CoV2 clusters inclusive of A2a, A3, and A4 were identified. A4 cluster was more prevalent in India ^12^. Comparison of the latest mutation data with these clusters revealed high frequency of mutations, such as nucleocapsid p.13P>L; nsp3 p.1197S>R; p.1198T>K; spike p.614D>G; RdRP (A97V), nsp2 p.159A>V; Orf3a p.57Q>H ^12^.

Among other viral variants, Δ382 reported in Singapore associated with lower odds of developing hypoxia requiring supplemental oxygen compared with infection with the wild-type virus ^24^. To prove that the viral genotype could affect infectivity, Yao et al, (2020) undertook functional characterization of mutations observed in their viral isolates. Very interestingly, these viral isolates showed significant variation in the cytopathic effects and viral load when checked by infecting Vero-E6 cells ^25^. In another study on data from public databases, Aiwewsakun et al, 2020 retrieved 152 full-length SARS-CoV-2 genomes from GISAID database, classified them into symptomatic and asymptomatic cases and evaluated the relationship between COVID-19 severity and virus genome variations ^10^. Nucleotide variations at the genomic position 11083 were associated with COVID-19 severity, with 11083G and 11083T observed more often in the symptomatic and asymptomatic patients, respectively. The authors also suggested that this could impact the interaction of host miRNAs with the viral genome, ultimately affecting the disease severity ^10^.

Apart from SARS-CoV-2 genome variations, host and host-agent interactions may impact the outcome of SARS-CoV-2 infection. The host factors may include, patients’ age, the degree of immune response to the virus due to difference in degree of T regulatory response (hygiene hypothesis), presence of co-morbid illness, Bacillus Calmette-Guerin (BCG) vaccination, and variation in the gut microbiota ^4–7^. In a well-designed lone GWAS study on host genetics, genetic susceptibility loci in COVID-19 patients with rs11385942 at locus 3p21.31 and rs657152 at locus 9q34.2 were found significant at the genome-wide level of significance (P<5×10−8). Apart from viral and host genome variations, a number of co-morbidities have been linked with poorer outcomes in COVID-19 patients ^26–28^. Of particular note are metabolic syndrome, cardiac diseases and respiratory pre-disposition, which adversely affect the eventual outcome in these patients ^29,30^. Another important factor that has been suggested to affect COVID-19 disease severity is the gut microbiome ^5^. Interestingly, a gut-lung axis has been suggested to influence the lung’s susceptibility to viral infections ^31^. We did not find a significant difference in the age of symptomatic patients from asymptomatic; nevertheless, the lack of adjustment of data for patient age, host genetic factors, microbiome and comorbidities is a limitation of our study.

In conclusion, the comparison of virus genotype across symptomatic and asymptomatic COVID-19 patients identified a clustering of unique viral genome variants in the symptomatic classes, which were completely absent in the asymptomatic class. However, these variants did not differ in frequency across the symptomatic classes showing no correlation with the degree of morbidity within the symptomatic class. Out of these variants, we would like to emphasize on the potential importance of the mutations observed in all three categories of the symptomatic patients. Among these g.241C>T mutation in the 5’UTR, p.1197S>R and p.1198T>K in the Nsp3 protein, p.97A>V in RdRp, p.614D>G and p.789Y>Y in the spike protein could be potentially important for disease severity. 5’UTR mutation g.241C>T could affect the stability of the transcript, and the synonymous mutation (p.789Y>Y) could affect translation by differences in codon usage preference in humans. Among other mutations, p.1197S>R and p.1198T>K in the Nsp3 protein, p.97A>V in RdRp and p.614D>G in the spike protein lie in crucial domains of the polymerase of the spike protein. D614G has already been reported to affect viral infectivity, and perhaps affects virulence as well. With available data, it appears that the 614G variant is a more infective form of the virus which causes less severe form of the disease in comparison to the 614D variant ^18,20^. We conclude that the virus genotype variations show a strong relationship with COVID-19 severity. This emphasizes a significant role of viral genome variations in COVID-19 symptoms, presentation and outcome.

## Acknowledgement

Singh Rajender would like to thank the Council of Scientific and Industrial Research (CSIR) for funding of SARS-CoV-2 sequencing under project code MLP2021. UCG thanks the Department of Biotechnology, Government of India, for funding (project No. BT/PR40311/COD/139/9/2020).

## Authors’ contributions

PM, SS, RV, UG, RM, RP, SR undertook sequencing work. SS, RV, AP, RS, DS undertook sample collection. UGC, UG, TKK, RR, SR planned the study. PM, SS, RM, RP, SR analyzed the sequence data. RR undertook structure-function analysis. PB undertook phylogenetic analysis. PM, RV, SR collected sequences data from other countries. PM, UG, UGC, RV, SS, TTK, SR wrote the article. All authors have read and agreed to this submission.

---

**Supplementary Table 1**. Relative synonymous codon usage patterns for SARS-CoV-2 genomes considered in this study.

**Supplementary Table 2**. Nucleotide frequencies for SARS-CoV-2 genomes considered in this study.

